# Modeling of Large-Scale Functional Brain Networks Based on Structural Connectivity from DTI: Comparison with EEG Derived Phase Coupling Networks and Evaluation of Alternative Methods along the Modeling Path

**DOI:** 10.1101/043109

**Authors:** Holger Finger, Marlene Bönstrup, Bastian Cheng, Arnaud Messé, Claus Hilgetag, Götz Thomalla, Christian Gerloff, Peter König

## Abstract

Here we use computational modeling of fast neural dynamics to explore the relationship between structural and functional coupling in a population of healthy subjects. We use DTI data to estimate structural connectivity and subsequently model phase couplings from band-limited oscillatory signals derived from multichannel EEG data. Our results show that about 23.4% of the variance in empirical networks of resting-state fast oscillations is explained by the underlying white matter architecture. By simulating functional connectivity using a simple reference model, the match between simulated and empirical functional connectivity further increases to 45.4%. In a second step, we use our modeling framework to explore several technical alternatives along the modeling path. First, we find that an augmentation of homotopic connections in the structural connectivity matrix improves the link to functional connectivity while a correction for fiber distance slightly decreases the performance of the model. Second, a more complex computational model based on Kuramoto oscillators leads to a slight improvement of the model fit. Third, we show that the comparison of modeled and empirical functional connectivity at source level is much more specific for the underlying structural connectivity. However, different source reconstruction algorithms gave comparable results. Of note, as the fourth finding, the model fit was much better if zero-phase lag components were preserved in the empirical functional connectome, indicating a considerable amount of functionally relevant synchrony taking place with near zero or zero-phase lag. The combination of the best performing alternatives at each stage in the pipeline results in a model that explains 54.4% of the variance in the empirical EEG functional connectivity. Our study shows that large-scale brain circuits of fast neural network synchrony strongly rely upon the structural connectome and simple computational models of neural activity can explain missing links in the structure-function relationship.

**Author Summary:** Brain imaging techniques are broadly divided into the two categories of structural and functional imaging. Structural imaging provides information about the static physical connectivity within the brain, while functional imaging provides data about the dynamic ongoing activation of brain areas. Computational models allow to bridge the gap between these two modalities and allow to gain new insights. Specifically, in this study, we use structural data from diffusion tractography recordings to model functional brain connectivity obtained from fast EEG dynamics. First, we present a simple reference procedure which consists of several steps to link the structural to the functional empirical data. Second, we systematically compare several alternative methods along the modeling path in order to assess their impact on the overall fit between simulations and empirical data. We explore preprocessing steps of the structural connectivity and different levels of complexity of the computational model. We highlight the importance of source reconstruction and compare commonly used source reconstruction algorithms and metrics to assess functional connectivity. Our results serve as an important orienting frame for the emerging field of brain network modeling.

## 1 Introduction

Resting-state brain activity represents the changes in neuroelectric or metabolic activity that occur when a subject is not performing a specific task and sensory input is largely reduced and stable. In this state spontaneous fluctuations emerge in the ongoing brain activity that synchronize across regions to exhibit a structured spatiotemporal pattern. Emerging resting-state networks have provided useful information regarding functional brain states, alterations in psychiatric or neurologic diseases, served as a basis for mapping and parceling the brain, and have helped to explain trial-to-trial fluctuations in cognitive functions [1, 2]. Although electrophysiological recordings of brain activity have already revealed ongoing activity a long time ago [3–5], the first description of common and organized networks emerging from ongoing activity was from functional Magnetic Resonance Imaging (fMRI)/Positron Emission Tomography (PET) studies which capture correlated slow fluctuations (< 0.1 Hz) across regions [6, 7]. Similarly, amplitude envelopes of alpha‐ and beta-frequency oscillations (~ 8 – 12 Hz and ~ 12 – 30 Hz respectively) display similar correlation patterns as the fMRI signals and are usually oscillating at a similar slow time scale of < 0.1 Hz. Both are here referred to as slow-fluctuating envelope resting-state networks.

The origin of resting-state ongoing brain activity is unresolved, but much evidence points to the anatomical skeleton shaping functional interactions between areas. Accordingly, relations of slowly oscillating resting-state networks (< 0.1 Hz) and long-range axonal connections have been detected, indicating that local activity of segregated brain regions is integrated by white matter pathways [8–10]. Although structural connectivity (SC) measured by diffusion tensor imaging (DTI) is a good predictor of functional connectivity (FC), functional connections also occur where there is little or no structural connectivity [8, 9]. Honey et al. found that some of the variance in FC that could not be related to structure could, however, be accounted for by indirect connections and interregional distance [9]. To explain missing links between anatomical structure and observed resting-state dynamics, bottom-up computational models based on structural priors offer interesting insights [8–10]. Different computational models reflecting various biological mechanisms for the emergence of the spatiotemporal dynamics of resting-state networks have helped to explain the variance between SC and spatiotemporally organized low-frequency fluctuations [11–14]. These dynamic simulations have robustly shown that the introduction of delays, scaling of coupling strength and well as additive noise lead to the emergence of functional patterns which resemble empirical resting-state networks operating in the low-frequency range.

Prior DTI-fMRI modeling studies have faced several technical challenges. First, the choice of computational model demands a trade off between highly simplified phenomenological models and biologically realistic models with high parameter space. Surprisingly, as shown by Messé et. al (2014), a simple stationary model of functional connectivity better explains functional connectivity than more complex models [15–17]. Second, preprocessing of DTI data is necessary to derive a structural connectivity matrix on a given parcellation scheme to overcome biases introduced by the latter. But the precise steps giving the most realistic structural connectome map are largely unknown.

Large-scale resting-state networks were originally described for correlated slow activity fluctuations recorded by fMRI/PET, or broadband power envelopes of the magneto-/electroencephalography (MEG/EEG) signal [18]. However, there is accumulating evidence that large-scale resting-state networks are also expressed in neuronal rhythms at faster frequencies [19, 20]. Fast fluctuations in neuroelectric activity, and especially the functional linkage of regions via phase correlations, are well known to underlie a broad variety of cognitive processes [21–24]. Synchronization of oscillatory neuronal activity among functionally specialized but widely distributed brain regions has been recognized as a major mechanism in the integration of sensory signals underlying perception and cognitive processes [25, 26].

Regarding the spatial organization of fast oscillatory phase correlations, its quantitative relationship to SC has not been investigated yet [27, 28]. Faster timescales of neural activity comprise for example the alpha, beta, or gamma band which constitute the major rhythms of spontaneous neuroelectric activity picked up by MEG/EEG. It has been argued, that compared to networks of slow fluctuations, structural connectivity does not strictly determine frequency-specific coupling in networks of ongoing activity at a faster timescale [28]. Indeed, phase coupling between segregated areas strongly relies on cortico-cortical connections [21, 29], implicating likewise a strong structure-function relationship.

### Performance of the reference model

In this study we probed this assumption of a strong structure-function relationship by simulating local node dynamics based on SC and comparing the phase relationships emerging from the simulated neural activity with empirically measured phase relationships. To this end, we combined SC from DTI data using probabilistic fiber tracking and FC from EEG data recorded during wakeful rest in 18 healthy individuals. We then used computational modeling approaches to link SC and empirical FC at fast frequencies. We demonstrate that empirical networks of resting-state fast oscillations are strongly determined by the underlying SC and that additional variance between structure and function can be explained by modeling dynamic activity based on white matter architecture. Specifically, the simulated FC explained 28.5% of the variance in the empirical FC that was left unexplained by SC alone. To further understand the explanatory power of our model we investigated its performance at the local level by assessing specific properties of ROIs (nodes) or connections (edges). We found that the model error was highest for large highly interacting ROIs.

However, modeling large-scale brain dynamics based on structural priors brings up several methodological alternatives, not only regarding the modeling itself, but also regarding alternative methods regarding the comparison of simulated and empirical data. Especially with resting-state MEG/EEG activity, the specificity of analytic routines requires methodological decisions which potentially lead to tremendous differences in modeling outcomes. We systematically assessed the effect of technical variations on results and their influence on the interpretation of structure-function relations. Specifically, we used our modeling framework to explore several technical alternatives along the modeling path and evaluate the alternative processing steps based on their effect on the performance of the model in simulating empirical FC. Specifically, we addressed the effects of five critical aspects in the modeling pipeline:

### Building the structural connectome

We used DTI and probabilistic tracking algorithms to compile a whole-brain structural connectome [30]. However, several studies suggested that current fiber tracking algorithms fail at capturing particularly transcallosal motor connections that are observed in non-human primate tracer studies [31, 32]. In addition, structural connection strength modeled by probabilistic tractography algorithms is influenced by fiber length due to the progressive dispersion of uncertainty along the fiber tract [33]. Therefore, we evaluateed the effect of normalizations for fiber length of the SC and examined the effect of weighting homotopic connections in our model. Our results show that the correction for fiber distance leads to a small decrease in the performance of our model. The additional weighting of homotopic transcallosal connections, however, increased the model fit [15, 16].

### Model of functional connectivity

Several alternative computational models of neural dynamics are available. In the choice of a more abstract version to a more realistic description of cortical interactions, these models vary in the complexity of their formulation and therefore might explain more or less variance in the observed FC. The downside of complex models, however, is the increased number of free parameters. These have to be approximated, need to be known a priori, or explored systematically. All these aproaches are problematic. For an assessment of the factor of model complexity, we compared a simple spatial autoregressive (SAR) model to the Kuramoto model of coupled oscillators. Surprisingly we find that the SAR model explains already a large portion of the variance and that the Kuramoto model only gives a slight improvement.

### Forward and inverse models

The comparatively few existing studies on large-scale modeling of MEG/EEG data differ systematically with respect to the comparison with empirical data. Some approaches project the observed time series onto the cortex using an inverse solution, whereas others project the simulated cortical signals into sensor space using the forward model [11, 34, 35]. We used our analytic framework to compare empirical and simulated FC at different spatial levels. We found that the importance of structural information is dramatically reduced, if the higher spatial resolution obtained by source reconstruction is bypassed.

### Source reconstruction algorithms

Estimating the spatiotemporal dynamics of neuronal currents in source space generating the EEG and MEG signals is an ill-posed problem, due to the vastly larger number of active sources compared to the number of sensors. Therefore, we assess the impact of specific source reconstruction algorithms on the match of simulated and empirical FC. We compared three routinely used algorithms that differ regarding the assumptions made about the source signal, such as smoothness, sparsity, norms, correlation between source signals. However, we found no compelling superiority of one algorithm over another.

### Functional connectivity metrics

Functional connectivity describes statistical dependencies between two signals often based on undirected temporal average such as correlation. In the last decades, various additional FC metrics have been introduced. These differ with regard to the relative weighting of phase and amplitude or concerning the removal of zero-phase lag components prior to correlation. The theoretical superiority of one approach over another is debated [36]. However, no consensus appears achieved and currently no single metric is dominantly used over the others. Therefore, we compared several widely used metrics to compare empirical and simulated FC. We found that the model fit was much better if zero-phase lag components were preserved in the empirical functional connectome.

In the following sections, we first present a reference procedure for modeling FC based on DTI and the comparison with empirical fast dynamics FC as measured by EEG. After an initial short overview of the modeling approach in section 2.1, we guide the reader step by step through the model details with the resulting outputs of each processing stage (section 2.2). From there, the impact of technical alternatives on the performance of the model is presented (section 2.3).

## 2 Results

### 2.1 Workflow

We compared the simulated FC based on SC with the empirical FC derived from EEG data (Figure 1). Our model includes the processing steps as shown in Figure 1 with the DTI measurements on the left and the EEG measurements on the right. We address preprocessing of DTI data in the form of homotopic reweighting. Then, the 66 ROIs of the cerebral cortex according to the ’Desikan-Killiany’ cortical atlas made available in the Freesurfer toolbox, were individually registered for 18 healthy subjects using Freesurfer (surfer.nmr.mgh.harvard.edu) [37]. The SAR model used in the reference procedure was selected based on simplicity and performance. We reconstructed source activity at the geometric center of each ROI based on the EEG time series by a linear constraint minimum variance spatial beam former (LCMV). Then we assessed FC between source time series band pass filtered at 8 Hz where the averaged coherence showed a peak (see supporting material S2 Frequencies). Finally, we evaluated the match of simulated and empirical FC based on the correlation between all pairs of ROIs [38]. Following this modeling approach, several alternative ways at each processing stage arise. Choices exist, for example, for the level of abstraction of the model type [39], metrics to compare functional connectivity and the approach to the inverse problem in interpreting EEG data.

**Figure 1.**
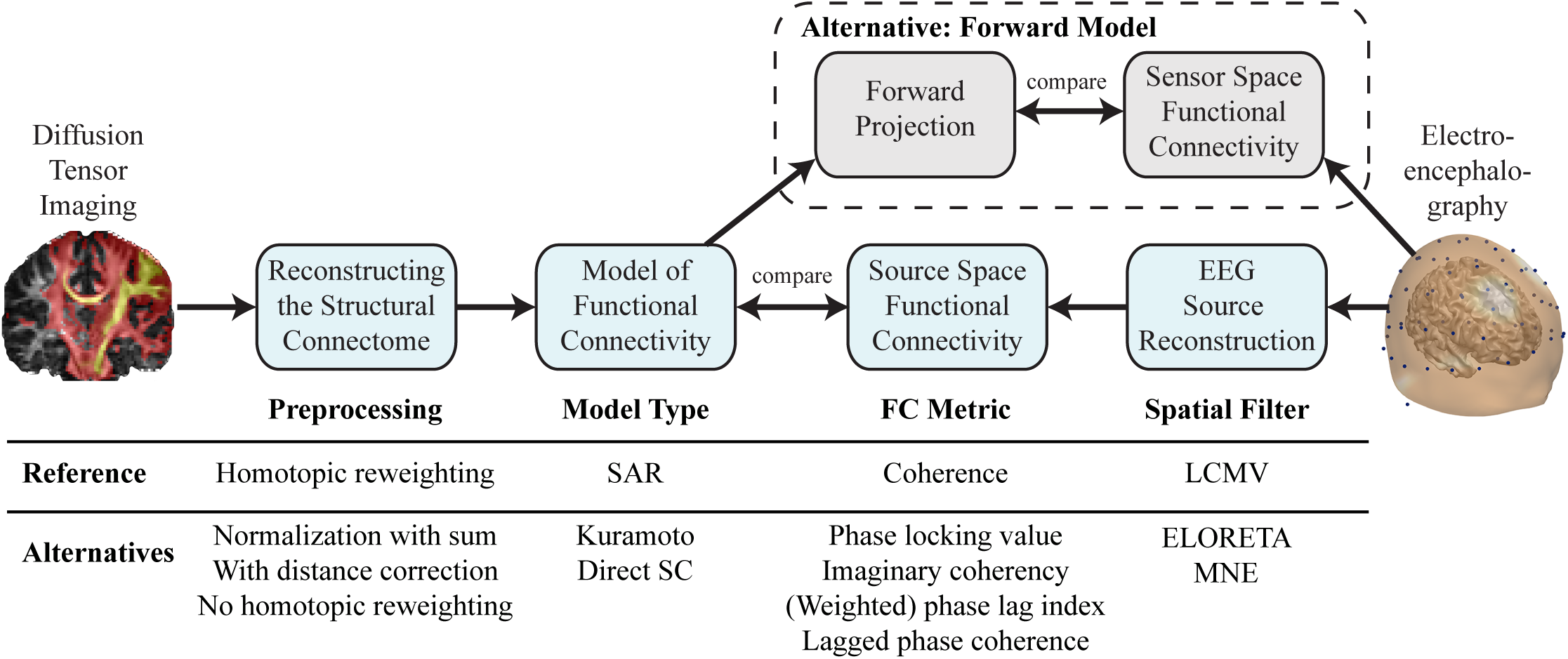
Workflow from DTI to the model of functional connectivity and comparison with empirical EEG data. Each processing step in the reference procedure can be replaced by several alternative methods. From left to right: Probabilistic tracts derived from DTI are preprocessed to give the structural connectivity matrix. From there we simulate functional connectivity and optimize free model parameters to maximize the global correlation with the empirical functional connectivity. The empirical functional connectivity is calculated between all pairs of ROIs after projecting EEG scalp recordings to source space using spatial filters. Alternatively, the comparison between simulated and empirical connectomes can be done in sensor space by projecting the simulated functional connectivity into sensor space using the leadfields.

### 2.2 Reference Procedure

#### Reconstructing the structural connectome

The assessment of individual SCs is based on the number of probabilistic fibers connecting the parcellated brain regions. In our reference procedure, four preprocessing steps were applied to the raw fiber counts: First, we normalized the total number of tracked fibers between two regions by the product of the size of both regions. This effectively normalizes the connection strength per unit volume [40]. Second, we excluded all self-connections by setting the diagonal elements of the SC matrix (denoted as S) to zero. The resulting SC matrix between the 66 anatomical ROIs is presented in Figure 2A. Previous studies showed that current fiber tracking algorithms underestimate transcallosal connectivity [31, 32]. Accordingly, modeling studies have revealed that specifically increasing the SC between homotopic regions leads to a general improvement of the predictive power irrespective of the model [15, 16]. Therefore, in the reference procedure we also increased the connection strength between homotopic regions by a fraction (h=0.1) of the original input strength at each node. Last, we normalized the input strength of each region to 1, as done in previous simulation studies [12, 15].

**Figure 2.**
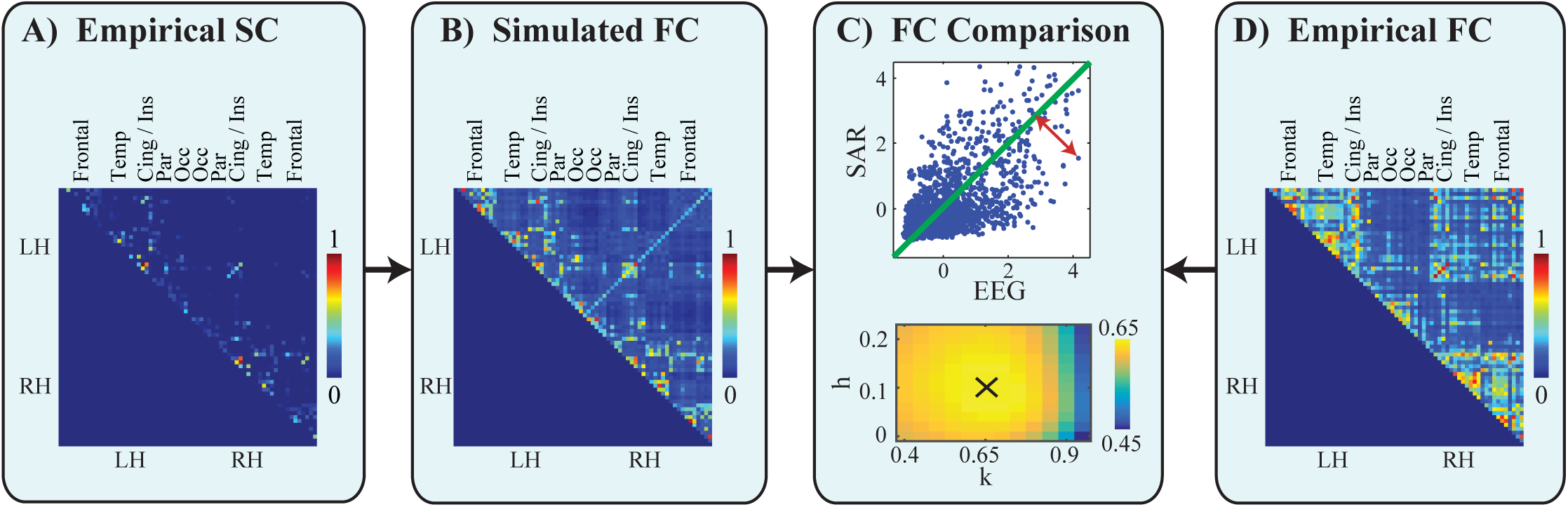
Comparison of empirical and simulated FC in the reference procedure. A: Structural connectivity among 66 cortical regions after normalization for ROI size and excluding self-connections (section 2.2, structural connectivity processing). B: The correlation of the simulated network based on structural connectivity using the SAR model with optimal global scaling parameter k=0.65 and homotopic connection strength h=0.1. C: Upper: The respective simulated (k=0.65, h=0.1) and empirical connection strengths are z-transformed and plotted for each connection. Correlation is used as a global performance measure. The local model error per connection is evaluated as the distance (red arrow) to the total-least-squares fit (green line). Lower: Color indicates the correlation strength at a range of different global connection strength scaling parameters k, and fraction of added homotopic connections (h). The black cross indicates the parameters with the maximum correlation. D: The empirical functional connectivity as the coherence between source reconstructed time series at the cortical regions. All connectivity matrices (A,B,D) were normalized to have strengths between 0 (no connection) and 1 (strong connection).

#### Model of functional connectivity

Several computational models of neural dynamics have been presented previously, varying in complexity regarding cellular and circuit properties [15, 41, 42]. In the reference procedure, we chose a model of FC which is as simple as possible while still explaining a substantial fraction of the variance in the empirical data. For resting-state FC derived from fMRI data, it was shown that the simple SAR model generates good matches at low computational expense [16, 17]. Therefore, we used the SAR model as a reference to evaluate just the static higher order dependencies in the FC.

The SAR model assumes that the time series of each region is a linear combination of the fluctuations of the time series of all other regions with added Gaussian noise, where only instantaneous effects are modelled. The activation of all ROIs 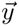 in the steady state condition is given by

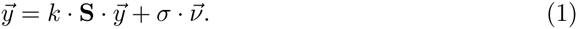

where **S** is set to the preprocessed SC matrix averaged across subjects as explained in the previous section. *k* is a global parameter describing the scaling of the coupling strengths. 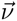 is uncorrelated Gaussian noise that is added at each node individually and is scaled by *σ*. This equation describes the equilibrium state of the autoregressive model.

The covariance between the time series of the SAR model can be solved analytically by substituting [43]

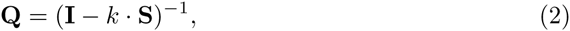

so that

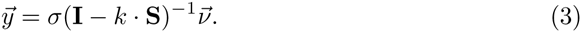

The covariance matrix between sources is then given by

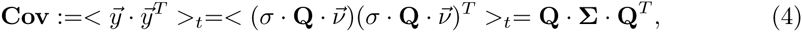

where <>_*t*_ denotes the average over time and 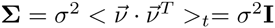 the noise covariance. Due to the assumption of uncorrelated Gaussian noise **Σ** is the identity matrix.

A FC is constructed based on all pairwise correlations between network nodes. This can be calculated using the standard definition of correlation given the covariance from equation 4:

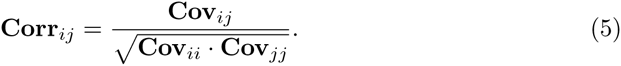

This step normalizes for different variances in the time series of different network nodes. The resulting correlation matrix, as shown in Figure 2B, is the predicted FC generated by the model given SC. The distribution of modeled FC is less sparse than the raw structural connection strength values: In SC (Figure 2A), many pairwise connections are close to zero and only few pairwise connections are large. To quantitatively evaluate the difference between the SC and the model output, we calculated the kurtosis of the values in the connectivity matrices:

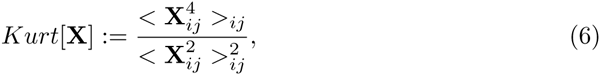

where <>_*ij*_ denotes the average over all upper triangular matrix elements without the diagonal (i.e. *i* < *j*). In this definition we divide the fourth raw moment by the second raw moment, where raw means that the moment is about the origin in contrast to central moments about the mean. The SC has a very high kurtosis (*Kurt*[*S*] = 62.83), whereas the FC predicted by the SAR model has a much smaller kurtosis (*Kurt*[*Corr*] = 5.77), indicating reduced sparsity.

#### Source reconstruction algorithms

The spatiotemporal dynamics of neuronal currents in source space can be estimated using various source reconstruction techniques applied to the MEG/EEG signal. The algorithms differ regarding the assumptions made about the source signal (i.e. smoothness, sparsity, norms, correlation between source signals). These assumptions about the signals to be reconstructed are a prerequisite to make the ill-posed inverse problem of distributed sources treatable. As a reference, we used a LCMV spatial beamformer, which reconstructs activity with unit gain under the constraint of minimizing temporal correlations between sources [44]. This approach has been applied in large-scale connectivity and global modeling studies before [11, 38, 45]. Multichannel EEG data was projected to source locations based on individual head models. The spatial filter was calculated for the optimal dipole orientation corresponding to the direction of maximum power, thus giving one time series per ROI. As a priori source locations we used the geometric center of each of the 66 ROIs individually registered on T1 images. See supplementary material for details on data acquisition, preprocessing and analysis of EEG data.

#### Functional connectivity metrics

FC can be assessed using several methodologies which differ with regard to the relative weighting of phase and amplitude or concerning the reduction of zero-phase lag components prior to correlation [46]. The choice of metric may have an influence on the match between empirical and simulated FC. In the reference procedure, we calculated ordinary coherence as a metric for FC due to its original and prepotent implementation in synchronization studies [25, 47–53]. The time series at each source were bandpass filtered at the alpha frequency range (8±2 Hz) and then Hilbert transformed. This choice of frequency was based on the general importance of the alpha rhythm for resting-state topographies [54, 55]. A broad spectrum (3-30 Hz) exploration showed a peak of the mean coherence across all connections at around 8 Hz (see supporting material S2 Frequencies).

The FC metrics are based on the analytic signal representation

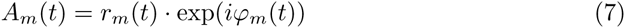

of region *m*. Furthermore, we calculated the cross-spectrum between two regions of interest m and n as

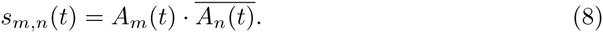

Given the analytic signal, the auto‐ and cross-spectra were computed and the coherence derived as the normalization of the cross-spectrum by the two auto-spectra [48]. This gives a FC index ranging from 0 to 1 between all pairs of ROIs:

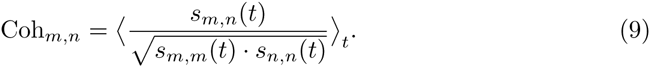

The resulting mean empirical FC matrix across the group is depicted in Figure 2D and was compared with the modeled FC matrix. Intrahemispherically, we found high connectivity within frontal and temporal areas in both hemispheres. Interhemispherically, the insular and cingulate areas were strongly connected.

#### Performance of the reference model

The SAR model yields a FC of the 66 parcellated brain regions in accordance with the empirical FC. Since both these matrices are symmetric, only the triangular parts are compared to assess the match between simulated and empirical FC. We calculate the performance of the model as the correlation between all modeled and empirical pairwise interactions Figure 2C. This performance metric is also commonly used in other studies [15, 41]. We found a high correlation between the FC from the model and EEG coherence values (r=0.674, n=2145, *p* < .0001) for the parameters k=0.65 (global parameter describing the scaling of the coupling strengths) and h=0.1 (additional weighting of the homotopic connections in the SC matrix) marked in Figure 2C below).

To put this into context, we first compared these results with the match between the empirical SC and FC without modeling (r=0.4833, n=2145, *p* < .0001) and found a shared variance of 23.4% (variance explained is 100 · *r*^2^). Modeling FC based on this SC backbone increased the global correlation to 45.4% (square of r=0.674). In other words, the modeled FC explains roughly 28.8% of the variance in the empirical FC that is left unexplained by SC alone.

To further understand the explanatory power of our model we investigate its performance at the local level by assessing specific properties of ROIs (nodes) or connections (edges). We defined for each connection the local model error as the distance (example shown as red arrow in Figure 2C, upper) between each dot and the total-least-squares fit (green line in Figure 2C, upper). Specifically, the question arises whether the high correlation between modeled and empirical FC is driven more by long or short edges. For example, the FC estimation between very close ROIs (in Euclidean space) might be spuriously inflated by volume conduction. Alternatively, there might be an overestimation of the SC between specifically close regions which could cause a higher model error [56]. To address this question we compared for each edge the model error with the fiber distance (Figure 3A). The average fiber distance between connected ROIs was negatively correlated with the logarithm of the local model error of each connection (r=-0.32, n=2145, *p* < .0001). A similar dependence was calculated between Euclidean distance between ROI locations and local model error (r=-0.33, n=2145, *p* < .0001). Both results indicate that the SAR model performed worse in simulating FC for closer ROIs in topographic space (measured in fiber lengths) and Euclidean space (measured as distance between ROI locations). This can be attributed to a higher variance in the SC and empirical FC matrices for close ROIs (as shown in the supporting material S3 Connection strength and distance).

**Figure 3.**
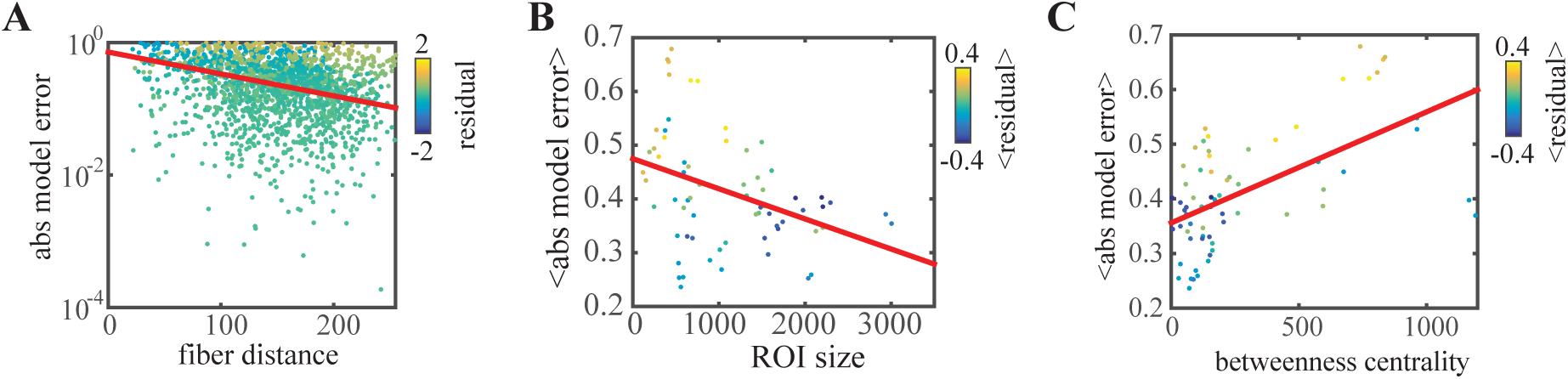
Dependence of residual and model error (absolute value of residual) on edge and node characteristics. A: linear fit of the log of the model error per connection showing a negative correlation with fiber distance. B: linear fit of the average model error per ROI showing a negative correlation with the size of the ROI. C: linear fit of the average model error per ROI showing a negative correlation with the betweenness centrality of the ROI. The angle brackets <> denote the average over all edges of the corresponding ROI. Residuals in A-C are calculated from the total least squares fit, negative values (blue dots) indicate that the average modeled functional connectivity per node was higher than the empirical functional connectivity, positive values (yellow dots) indicate that the the modeled functional connectivity per node was smaller than the empirical functional connectivity.

We further evaluated the performance in relation to certain node characteristics and averaged the errors of all edges per node. The node performance in terms of model error is shown in Figure 3B-D dependent on different node characteristics. First, we looked at the influence of ROI size on the model error. We hypothesized that due to larger sample sizes and more precise localization, the model error would be smaller for large ROIs. As expected, the model error for each ROI is negatively correlated with the corresponding size of the ROI (r=-0.37, n=66, *p* < .005) as shown in Figure 3B. Then we hypothesized, that due to the sparseness of SC, some ROIs in SC have a very high connectedness compared to functional data, leading to a larger model error. To address this aspect we calculated several graph theoretical measures that assess the local connectedness in different ways and related this to the average model error. As a first measure we calculated for each node the betweenness centrality, defined as the fraction of all shortest paths in the network that pass through a given node [57]. The absolute model error is positively correlated with the betweenness centrality (*r* = 0.58, *n* = 66, *p* < .0001) as shown in Figure 3C. A similar indicator of a nodes connectedness in the network is the sum of all connection strengths of that node. Also for this metric, we find a linear relationship between the total connection strength of a node and the model error (*r* = 0.35, *n* = 66, *p* < .005). In addition, the dependence between the model error and the eigenvalue centrality, which measures how well a node is linked to other network nodes [58], was evaluated (r=0.26, n=66, *p* < .05). The local clustering coefficient, which quantifies how frequently the neighbors of one node are neighbors to each other [59], did not show significant relations with the local model error (r=0.06, n=66, *p* = .65).

Overall, the reference model can explain much of the variance in the empricial FC. The error in the predicted FC of the reference model appears to be highest for small highly interacting ROIs. This might be due to the more heterogeneous structure of small highly interacting ROIs. On the other side, interactions between more distant and large ROIs are better predicted by the model, probably due to the more homogenous connectivity.

### 2.3 Alternative modeling approaches

The modeling of large-scale brain dynamics based on structural priors brings up several methodological alternatives. As a principal choice, the model may be evaluated either in source or in sensor space. In the baseline model that was presented above, we made specific choices at each processing stage based on simplicity and good explanatory performance. Especially with resting-state EEG activity, a lack of analytic routines requires methodological decisions to be made heuristically, which could potentially lead to substantial differences in the conclusions drawn. In the following section we systematically compare different alternatives of the procedural stages delineated above and compare the outcome regarding global correlation between simulated and empirical FC. First, we assessed the influence of distance normalization and weighting of homotopic connections in the structural connectome on simulated FC. Second, we tested if a more complex simulation model of coupled oscillators is able to capture a larger part of the variance of the empirical data that is not explained the simple SAR model. Third, we evaluated an alternative comparison in the sensor space using a forward projection of the source time series in contrast to source reconstruction. Then, we compared different source reconstruction methods. Finally, we tested the impact of removing zero-phase lags in functional interactions.

#### Reconstructing the structural connectome

The structural connectome was compiled using global probabilistic tractography. Interregional connections (edges) of the brain are represented by the number of ”probabilistic streamlines” between these regions (nodes). We tested the performance of two alternative modifications of the SC (Figure 4).

**Figure 4.**
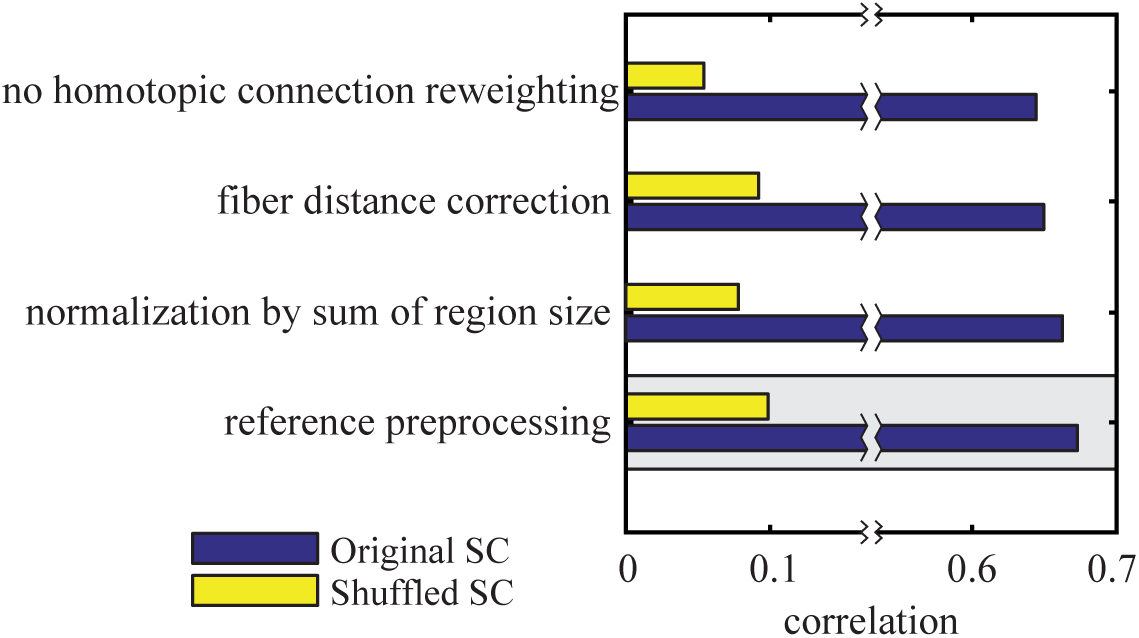
Structural connectivity preprocessing. The correlation between modeled and empirical functional connectivity for different preprocessing steps of structural connectivity. In the reference procedure, the number of tracked fibers between two regions was normalized by the product of the region sizes. The model based on the original structural connectivity is shown in blue and the baseline model which is based on shuffled structural connectivity in yellow. The gray box marks the reference procedure.

The pooled connectivity results obtained by the probabilistic fiber tracking are directly proportional to the size of the seed and target regions. The size of the regions, determined by the parcellation scheme, vary [9]. They are parcelated based on standard gyral-based neuroanatomical regions [37]. In order to account for a bias of stronger connectivity of larger regions, SC was normalized using the size of the regions. However, the exact method of normalization for ROI size is currently a matter of debate and no operational routine has emerged yet [60]. Therefore, we compared different normalizations regarding the quality of the model. In the reference procedure, we normalized the number of tracked fibers between two regions by the product of the region sizes. We found that this approach gives the best model performance (r=0.674, n=2145, *p* < .0001) in comparison with alternative normalizations that are presented in the following paragraphs.

First, instead of the normalization by the product of the two ROI sizes it is possible to normalize using the sum [15]. However, the performance decrease in comparison to the reference procedure is very small (r = 0.65, n = 2145, *p* < .0001).

Second, an additional weighting was applied to correct for the influence of fiber length on the probabilistic tracking algorithm. Therefore, the the number of streamlines connecting two regions was multiplied by the average fiber length between these areas. This normalization leads to a small decrease in performance (*r* = 0.64, *n* = 2145, *p* < .0001).

Third, we tested the influence of homotopic transcallosal connections by omitting the additional weighting applied in the reference procedure. As a result, the correlation between modeled and empirical FC drops from *r* = 0.674 to *r* = 0.63. These results demonstrate that our reference method of reconstructing SC is slightly superior to the evaluated alternative approaches. Overall, the performance of the empirical simulation based on the SC is rather robust with respect to the choices of preprocessing.

#### Model of functional connectivity

In the previous sections we showed that a considerable amount of variance in empirical FC can be explained even with a simple SAR model that captures only stationary dynamics. Several alternative computational models of neural dynamics have been presented that vary regarding their complexity. More complex models can incorporate aspects of cortical processing at the microscopic scale such as cellular subpopulations with differing membrane characteristics or, at the macroscopic scale, time delays between nodes [39, 41, 61]. The downside of complex models is the increased number of free parameters whose values need to be approximated, have to be known a priori, or explored systematically. We hypothesized that a more complex model which incorporates more parameters in order to simulate neural dynamics more realistically might explain more variance in FC. We decided to use the Kuramoto model of coupled oscillators as an alternative to investigate whether this holds true [62, 63]. In contrast to the SAR model, the Kuramoto model can incorporate delays between nodes and thus becomes a model of dynamic neural processes [42, 64]. At the same time the Kuramoto model is simple enough to systematically explore the parameter space. The progression of the phase of each neuron is modeled by the differential equation

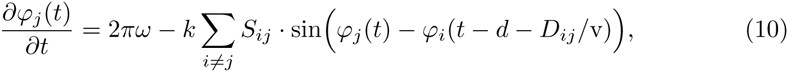

where *d* is a fixed delay at each node and *v* is the transmission velocity which is multiplied by the distance *D*_*ij*_, (see S1 Empirical Data) which leads to a connection-specific delay. The Kuramoto model was simulated using the Euler integration method in time steps of 0.1 ms. In contrast to the SAR model, which does not reflect temporal dynamics, in the Kuramoto model we used the same bandpass filters and coherence estimation method as described in equations 7, 8, and 9.

An additional alternative to the SAR model is an even more simple direct comparison between the empirical SC and FC. The simple structure-function comparison gave a 23.4% match between structural and functional connectivity alone (r=0.4833, n=2145, *p* < .0001). The SAR model and the Kuramoto model both explain more variance of the functional connectivity than this direct comparison of structural and functional connectivity (Figure 5A). Using the SAR model we simulated a functional connectome with a 45.4% match to the empirical data (r=0.674, n=2145, *p* < .0001). With the Kuramoto model however, the match could be further increased to 54.0% (r=0.735, n=2145, *p* < .0001). In other words, the modeled FC using the Kuramoto model explains 40.0% of the variance in the empirical functional connectivity that is unexplained by structure alone. In addition, demonstrating the importance of the underlying structural network, all three variants have a significantly higher correlation than for the randomly shuffled SC.

**Figure 5.**
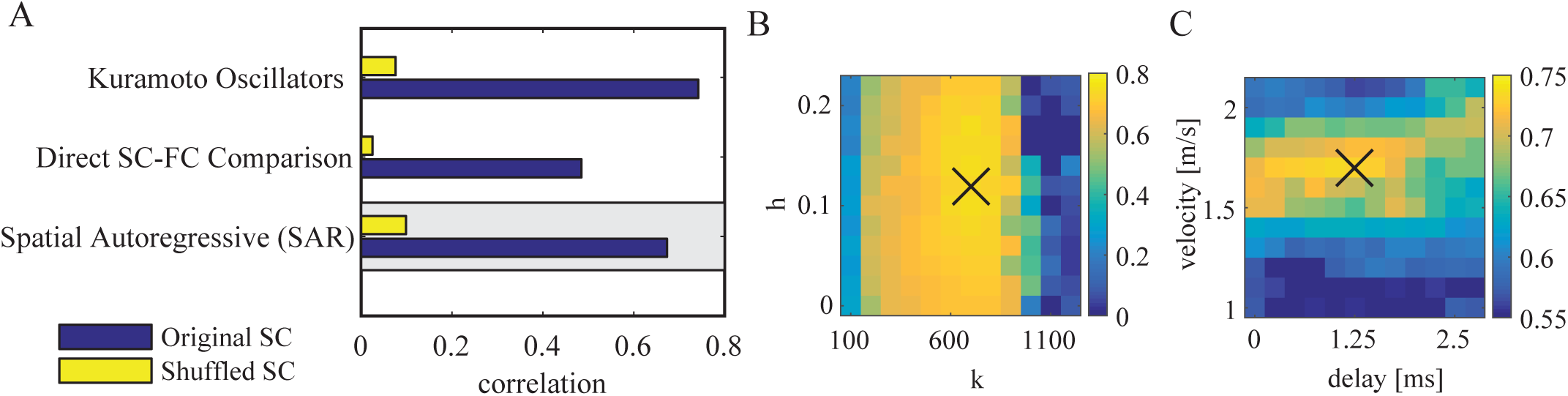
Model of functional connectivity. A: Performance comparison between the SAR model (reference model), the Kuramoto model and directly between the empirical and structural connectivity. The model based on the original structural connectivity is shown in blue and the baseline model which is based on shuffled structural connectivity in yellow. The gray box marks the reference procedure based on the SAR model. B: Performance of the Kuramoto model for different parameters k and h close to the optimal point with fixed velocity = 1.7 m/s and delay = 1.25 ms. C: Same as B but with varying velocity *v* and delay *d* with fixed *k* = 700 and h = 0.12. In panels B and C the X marks the parameter that was selected for the corresponding other panel.

The Kuramoto model showed the best performance for a connection strength scaling of *k* = 700 (Figure 5B). Important to note is that the constant delay can be neglected without a large performance drop (Figure 5C). In contrast, the velocity introduces a connection specific delay that is modulated by the DTI fiber lengths and the model performance has a considerable peak around *v* ≈ 1.7.

#### Forward and inverse models

In the comparatively few studies on large-scale modeling of MEG/EEG data, a discrepancy exists to whether simulations are compared with empirical data in the source or sensor space [11, 34, 35]. In other words, the measured time series are either projected onto the cortex using an inverse solution or the simulated cortical signals are projected into sensor space using a forward model. Here we compare both approaches, source reconstruction vs. forward projection, with respect to the global correlation strength between modeled and empirical FC. The source reconstruction approach has been described above (see section 2.2 and S1 Empirical Data).

For the inverse solution and forward projection, we computed as a forward model a boundary element method volume conduction model based on individual T1-weighted structural MRI of the whole brain and comprising 8196 dipoles distributed over 66 regions [65]. Each dipole has six degrees of freedom defining its position, orientation, and strength in the cortex. The positions for each vertex are defined to be lying equally spaced within the parcellated brain regions of the cortical sheet. The electric source activity can be approximated by the fluctuation of equivalent current dipoles generated by excitatory neurons that have dendritic trees oriented perpendicular to the cortical surface [34]. For the inverse solution, the dipoles orientation was assessed according to its maximal power. For the forward projection of simulated time series, the dipole orientations were defined by the normal vector of the cortical surface of the corresponding region in the segmented MRI image. Since each of the parcellated brain regions extends over several surface vertices, all dipole normals within each region are averaged. This results in one average direction vector per region (average length over all regions: 0.52) which is used to project into the EEG sensor space.

In the previous sections we showed that the underlying SC had a large impact on the relatively good match between simulated and empirical FC. Figure 4 and Figure 5A show large drops in correlation when the simulation is based on shuffled SC (yellow bars) instead of the original SC (blue bars). By comparing the source reconstruction with the forward model approach, we find that the comparison in sensor space using the forward projection yields higher correlations between simulated and empirical data (Figure 6A). If, however, the underlying structural connectivity is shuffled before applying the SAR model, the correlation of simulated and empirical FC remains equally high in sensor space. This indicates that the importance of structural information is dramatically reduced if the higher spatial resolution obtained by source reconstruction is bypassed. The forward projection of the simulated time series leads to a very low spatial specificity of the functional connectivities in sensor space (Figure 6B).

**Figure 6.**
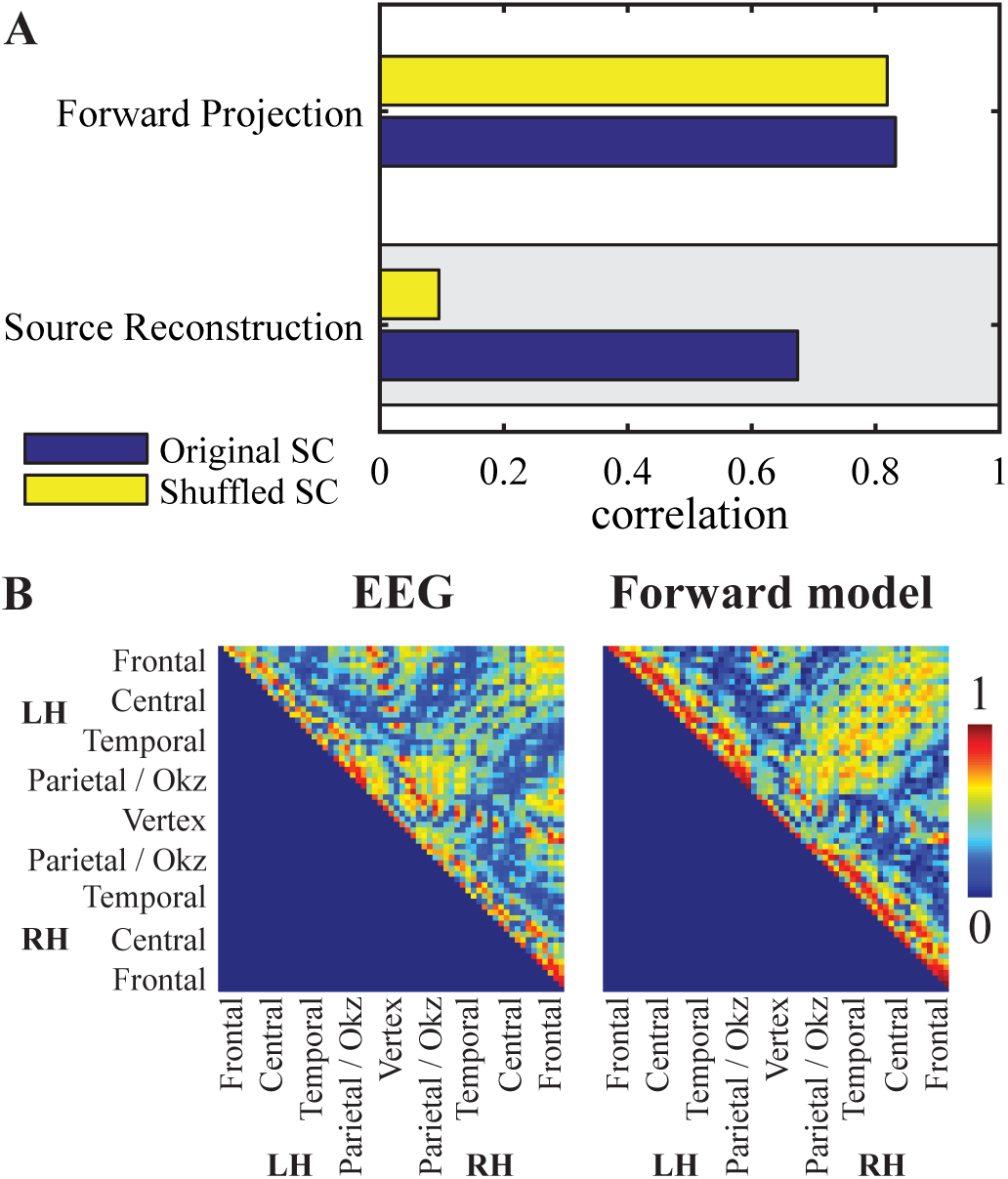
Comparisons of forward projection and source reconstruction. A: Global correlation between simulated and empirical functional connectivity in sensor space by applying the forward projection to the SAR model, or in source space by applying the LCMV beamformer to the EEG time series. Blue bars show simulations based on original structural connectivity and yellow bars simulations for randomly shuffled structural connectivity. The gray box marks the reference procedure. B: EEG functional connectivity measured by coherence (left) and the forward projected modeled functional connectivity (right), both in sensor space.

Since several inverse methods are routinely used without a clear superiority of one over another, we aimed to assess the impact of the specific source reconstruction algorithm on the fit between simulated and empirical FC. We compared three prominent and widely used inverse methods which make fundamentally different assumptions. (Figure 7). As a reference, we used an LCMV spatial beamformer which reconstructs activity with the constraint of minimizing temporal correlations between sources [44]. For comparison we calculated the inverse solution by using exact low resolution brain electromagnetic tomography (ELORETA) which reconstructs activity by spatial smoothness constraints and in this sense it emphasizes local temporal correlations in comparison to beamforming approaches [66]. It is also widely used in source connectivity analyses [67, 68]. Additionally we calculated the minimum-norm estimate (MNE) which recovers source activity by reducing overall energy [69] which is based on the assumption that the data gives no information about the null space component of the leadfield which is thus set to zero. Figure 7 shows the global correlation values resulting from these three alternative inverse solutions. It can be seen that all of them have a similar performance level (LCMV:r=0.674, n=2145, *p* < .0001), ELORETA: (r=0.728, n=2145, *p* < .0001), MNE: (r=0.676, n=2145, *p* < .0001). The connectivity maps of time series of the inverse solutions were highly correlations (LCMV-ELORETA: r=0.84, LCMV-MNE: r=0.95,MNE-ELORTEA: r=0.84; all *p* < .0001).

**Figure 7.**
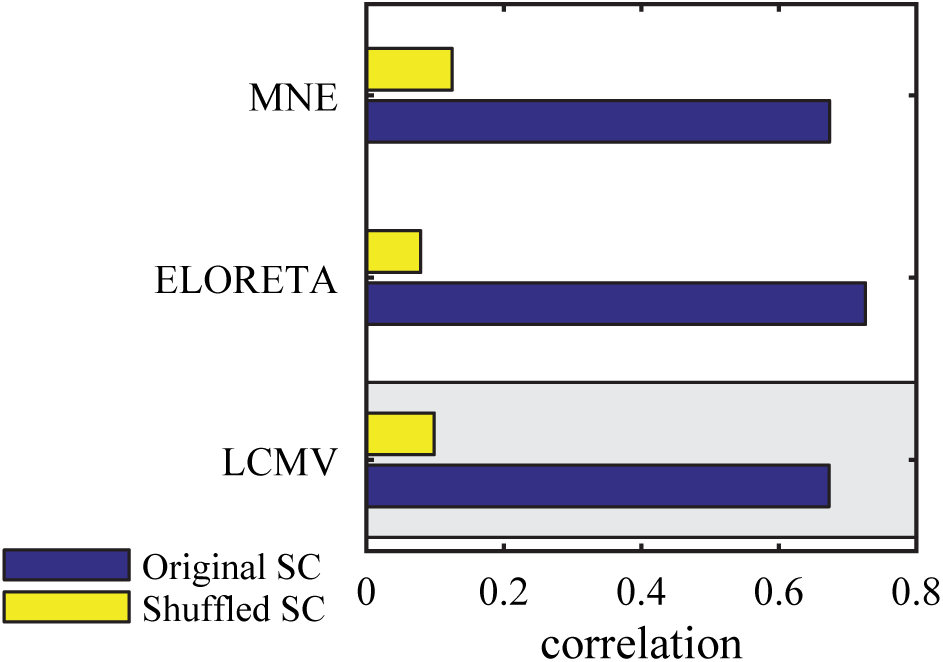
Source reconstruction. The correlation between modeled and empirical functional connectivity for different source reconstruction algorithms. The model based on the original structural connectivity is shown in blue and the baseline model which is based on shuffled structural connectivity in yellow. The gray box marks the reference procedure.

#### Functional connectivity metrics

We compared several widely used FC metrics regarding the global relation between empirical and simulated functional connectivity. Previous modeling studies implemented different metrics, and clear superiority of one over another has not been shown [36, 46, 70]. In the reference procedure, empirical FC was calculated as ordinary coherence and compared to the FC matrix derived from the SAR model. In addition, we investigated several alternative FC metrics.

All metrics were based on the same analytic signal representation as shown in equation 7 and the cross-spectrum as defined in equation 8. The different metrics are listed in Table 1 with their corresponding equations, characteristics and results. Comparing the performances based on all five measures (see Figure 8), we found a high correspondence in model performance between coherence and PLV. In contrast, PLI, WPLI, and LPC all showed a significantly lower match between simulated and empirical FC, with correlation coefficients between 0.10 and 0.18. ICOH showed the smallest correlation between modeled and empirical data with a non-significant p-value (r=0.103, n=2145, p=.37). For all metrics, the global correlation essentially vanished if the underlying SC was shuffled prior to simulation.

**Table 1.**
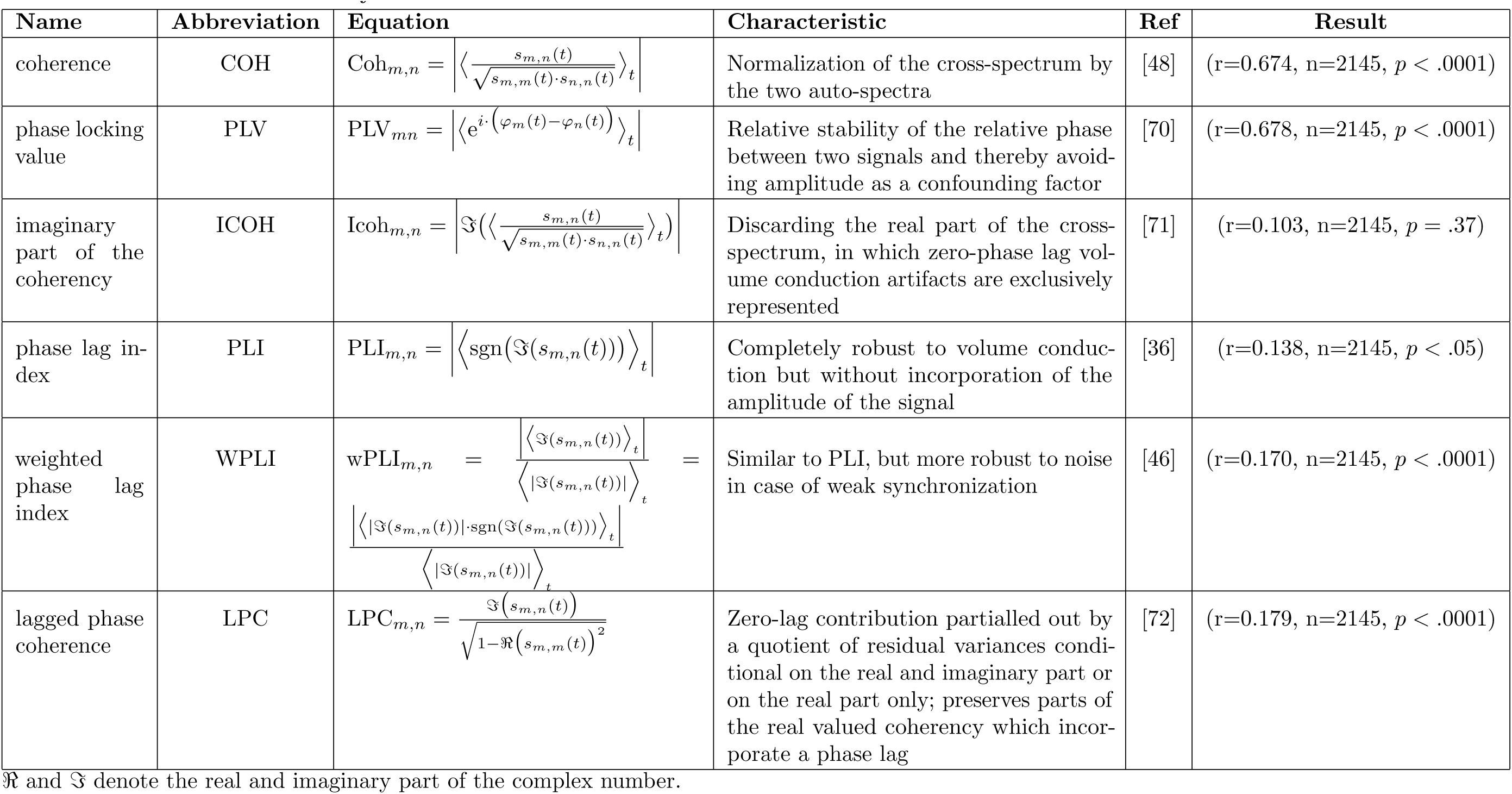
Functional Connectivity Metrics

**Figure 8.**
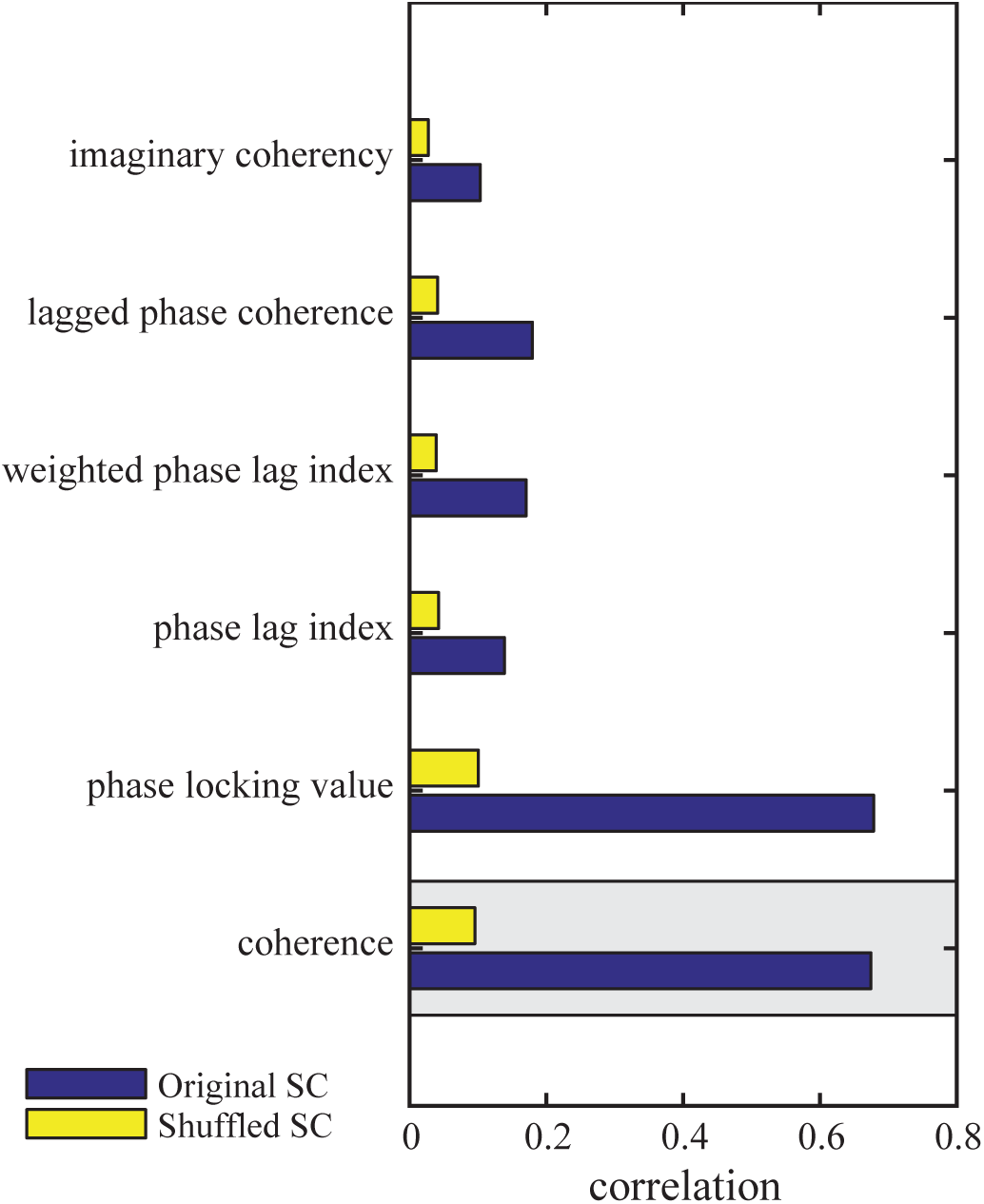
Functional connectivity metrics. The bars show the correlation between the empirical functional connectivity and the simulated functional connectivity obtained using the SAR model. The model based on the original structural connectivity is shown in blue and the baseline model which is based on shuffled structural connectivity in yellow. The gray box marks the reference procedure.

In summary, there are substantial decisions to be made at each stage of the processing pipeline. We selected the reference procedure prior to the evaluation of all alternatives in all these processing stages. Then, we evaluated the combination of all best performing alternatives along the pipeline. This best performing combination consists of the reference preprocessing of DTI data to construct SC, the Kuramoto oscillator network to simulate FC, PLV as a FC metric, and ELORETA as source reconstruction method from EEG. This combination results in a match of 54.4% between simulated and empirical functional connectivity (r=0.7377, n=2145, *p* < .0001).

## 3 Discussion

With this study we contribute to resolving the structure-function relationship in global connectomics. We simulated fast neural dynamics based on a realistic structural connectivity backbone and compare it to empirical functional connectivity derived from phase coupling of oscillatory brain waves. For the empirical data collected in 18 subjects we found a 23.4% match between structural and functional connectivity alone. Using a simple SAR model to simulate FC based on SC, this match was increased to 45.4%, showing that the model can capture about 28.8% of the variance in the empirical FC that is unexplained by the structure alone. We demonstrate several technical alternatives in the modeling procedure and derivation of empirical connectomes that are commonly used, but only few gave noticeable improvements. Of note, introducing additional model parameters by using the Kuramoto model of coupled oscillators improved the simulation (Figure 5). Our results show that resting-state networks emerging from phase coupling at a fast timescale largely resembles structural connectivity, as it has been previously shown for slow fluctuations of BOLD-signal or broad-band power envelopes [8, 9, 11, 13].

### 3.1 Modeling fast dynamics

It has been assumed that for the resting-state networks based on fast dynamics the underlying anatomical skeleton is less important compared to the slow resting-state networks, but this issue has not yet been systematically investigated [28]. We calculated the performance of the reference model as the correlation between all modeled pairwise interactions and all empirical pairwise interactions in an empirical functional phase relation connectome of the alpha rhythm and found a good match of 45.4% (Figure 2C). This finding is in contrast to the prior assumptions and shows that the anatomical skeleton is equally crucial for fast timescale functional interactions [21, 29].

To better understand the reference model performance we investigated the model error in relation to node and edge characteristics (Figure 3). In general, the model error decreased with longer fiber distance and Euclidean distance. Specifically, for short fiber distances, the model overestimated FC (negative residuals blue in Figure 3). Why are short connections in general more difficult to model based on white matter tracts? The empirical connectome was extracted from resting-state alpha topographies in which propagating waves play an important role for adjacent and remote brain areas to communicate with each other. Cortico-cortical axons in the white matter tracts are considered as the major route for traveling waves. However, a recent study presented compelling evidence for intracortical axons accounting for spatial propagation of alpha oscillations [73]. Such a mechanism would enable high local synchrony in the relative absence of structural connectivity measured by DTI.

We used a stepwise linear model to extract node characteristics explaining most of the model error and found that ROI size and betweenness centrality play an important role. Regarding ROI size, the smaller model error for larger ROIs could be attributed to the measurements of structural as well as functional connectivity being more reliable for larger ROIs: In the case of the SC measurements using DTI, a larger parcellated cortical region allows to track more streamlines with different initial conditions (i.e. for more voxels) and thereby allows a more reliable estimation of the connection probabilities between regions. In EEG as well as DTI, the localization and inter-subject registration of large ROIs can be assumed to be less effected by small deviations because a small spatial shift of a large ROI still allows a large overlap with the correct ROI volume whereas a small spatial shift of a small ROI could displace it completely outside of the original volume. For betweenness centrality, the opposite scenario was the case: the smaller the betweenness centrality the smaller was the model error. Central hubs in a structural network offer anatomical bridges which enable functional links between regions that are structurally not directly related [57]. Hard-wired connections do not necessarily contribute at all times to FC in the network and, vice-versa, functionally relevant connections do not necessarily have to be strongly hard-wired [9]. Possibly, the simple SAR model, which captures only stationary dynamics, has weaknesses at these central hub nodes. In order to capture the empirical FC at these nodes, a more complex dynamical model able to capture non-stationary dynamics with context switches at slower time scales is needed. Nodes with a high betweenness centrality can be expected to communicate with certain cortical modules only at certain times in specific dynamical regimes. We hypothesize that a more complex dynamical model of neural activity could capture this behavior more accurately. Therefore we suggest that further research could especially improve the model in these cases of dynamical context switches in central hub nodes, which cannot be captured by simple models such as the SAR model.

### 3.2 Reconstructing the structural connectome

Using our modeling framework to compare different alternatives of reconstruction the structural connectome, we found that the best match between simulated and empirical FC was obtained when an additional weighting of connections between homotopic transcallosal regions was applied. Additional weighting for fiber distances did not improve the simulation performance significantly. Overall, the differences were very small proving the modeling approach to be rather robust regarding the evaluated choices of reconstruction.

Currently, there is no common approach to correct for the influence of fiber distance on the probabilistic tracking algorithm [13, 33, 74]. Although we found that the model error was largest for small fiber distances (modeled FC higher than empirical FC), a correction for fiber lengths did not improve the result of the simulation. This suggests that the high local connection strength of SC obtained by DTI reflects actual structural connectivity. Methodically, this finding is supported by the fact that accuracy of probabilistic fiber reconstrunction decreases with distance, whereas short-distance connections are reconstructed with high reliability [31]. However, it remains a challenge to correct probabilistic tracking results for the impact of fiber distance and further work is needed to address this methodological limitation.

Our model improved with an additional added weight of homotopic connections, which is supporting the data by Messé et al. [15]. This finding points to a related limitation of the probabilistic tracking algorithms to correctly assess long distance and lateral transcallosal fibers. In agreement with previous studies, we show that this limitation can be addressed by adding an preprocessing step to the structural connectome reconstruction.

### 3.3 Model of neural activity

We show that our SAR model already explains much of the variance in the empirical EEG data. Our results indicate that the Kuramoto model moderately improved results compared to the reference model. The SAR model has a small number of parameters allowing a fast exploration of the parameter space [43]. The SAR model served several studies in which complexity and information-theoretical measures characterizing FC were explored [43, 75, 76]. As a downside, the SAR model has a smaller number of parameters and therefore lacks the modeling capacity to further optimize the dynamics to better fit to the empirical data. Furthermore, the SAR model cannot model individual frequencies and their interactions, making the Kuramoto model a viable alternative. It has been shown that the Kuramoto model features complex synchronization dynamics which can be related to the explanation of oscillatory phenomena in the human cortex, such as fluctuating beta oscillations [42] or metastable synchronization states [11]. A more detailed analysis of the synchronization properties of the Kuramoto model in the human connectome was done by Villegas et al. [77], where frustration and the transition between synchronous and asynchronous phases were analyzed [78]. The Kuramoto model was also used to study the effects of lesions on cortical dynamics and binding by synchrony [63, 79]. However, it has been shown that more complex models with more parameters are usually not better in explaining fMRI functional connectivity from structural data [15–17]. Highly parameterized models which require the numerical integration of differential equations take several orders of magnitude more computational time to obtain a reliable estimate of FC than the simple model used here. For certain neurophysiological questions however, the wider parameter space of complex models can be used to explore neural processing properties. The relative benefit of a dynamical model has to counterbalance the higher computational demand. Therefore, the choice of model depends on the investigated scientific question [17, 39]. In this study we used the simpler SAR model as a reference because the focus was to investigate alternatives also in many other stages of the processing pipeline and a more complex simulation model would impede identifying the best alternative in the other stages of the processing pipeline, due to the high dimensional parameter space.

### 3.4 Source reconstruction versus forward projection

The inverse problem is ill-posed since the higher number of possible active neuronal sources is higher than the number of recording channels. Thus, the ground truth of brain activity patterns generating the measured signal is impossible to infer. A variety of alternative methodological approaches have been developed regarding source imaging.

Particular caution should be exercised concerning the influence of different inverse solutions on the resulting data [80]. Here, we presented a comparison of the performance of three commonly used inverse methods regarding the global correlation between empirical and simulated FC in our technical framework (Figure 7). All source reconstruction algorithms perform in a similar range with resembling r between 0.674 and 0.728. Although the algorithms differ regarding the assumptions made about the source signal, the high correspondence in performance of the three source reconstruction techniques mutually validates their respective inverse solutions.

Next, we aimed to investigate the best approach for comparing empirical and simulated FC particularly in sensor and source space, see Figure 1. In the sensor space scenario, the simulated signal, as the mean field source activity generated by the Kuramoto model, was projected into sensor space to generate a simulated EEG signal by applying the leadfield (i.e. forward model). For this approach we found slightly higher correlations between simulated and empirical data (Figure 6A). However, we also found that the high correspondence between empirical and simulated EEG sensor space FC was independent of the underlying SC: Shuffling SC before the simulation did not abolish the correlation between the empirical and simulated FC as was the case when the comparison was done at the source space level. This lack of specificity of the simulated FC regarding the anatomical skeleton strongly suggests that the sensor level connectivity matrix is shaped mainly by the leadfield (Figure 6). In fact, the leadfield and can already explain most of the variance (81.9%) in the empirical FC of the sensor space. In contrast, the inverse solution in the source reconstruction procedure removes much of these volume conduction correlations so that the comparison of coherence in source space appears reasonable. We conclude that the volume conduction model of the head is mixing the source time series such that the coherence in sensor space reflects to a high degree the structure within this mixing matrix and the sensor space is a suboptimal stage for investigating structure-function relationships by large-scale modeling approaches. Thus, one should refrain from such a comparison in sensor space with metrics that do not exclude zero-lag interactions. In order to assess the accuracy of simulated global network characteristics, the comparative spatial level should be at source space in order to avoid signal mixing by the leadfield matrix and allow to include zero-lag interactions. The results offer an important ground for modeling studies using source connectivity analyses for MEG/EEG data.

### 3.5 Connectivity metrics and the role of phase lags

One of the main differences between fMRI/PET and MEG/EEG connectivity studies is that for MEG/EEG a multitude of different metrics to quantify FC are currently available and no single metric is predominantly employed or has emerged as being superior [81, 82]. This issue hampers comparability between studies and physiologic interpretation. It was our aim to use our theoretic framework for a systematic comparison of different functional connectivity metrics. We compared six commonly used metrics that differ regarding their sensitivity towards zero-phase lag coupling and amplitude variations. The definition of PLV, PLI, WPLI and LPC theoretically renders those metrics insensitive to amplitude variations. We found no major difference in performance between COH and PLV and no major difference between ICOH, PLI, WPLI, and LPC. This result is easily understood on the basis that the SAR model presents the steady state solution including a small noise component only. An important finding is the high correspondence in model performance between coherence and PLV. Coherence is the cross-spectrum between two sensors normalized with the auto-spectra whereas PLV quantifies the consistency of a phase difference between two signals across time. Both measures are high if there is a consistent phase difference regardless of whether the latter is near zero, 180° or inbetween. Similar results between coherence and PLV have been found in previous studies [83–85]. The similarity of both measures in our study suggests that amplitude variations between areas are of less weight than phase variations. Another main finding is the drop in model performance with the metrics ICOH, PLI, WPLI and LPC which are by design less sensitive to zero-phase coupling. Regarding the latter, a major concern exists whether such coupling in scalp recordings would be contaminated by volume conduction artifacts. Obviously, synchrony at sensor level could result from two channels picking up activity from a common source since the activity of the source signal passes through the layers of cerebrospinal fluid, dura, scalp and skull acting as a spatial filter. This effect leads to the detection of spurious synchrony, even if the underlying sources are independent [86]. Based on the assumption that the quasi-static approximation holds true for EEG, volume conduction would occur with zero-phase lag [87]. Thus, the most commonly used approach to deal with the problem of volume conduction is to neglect interactions that have no phase delay. This is, however, a potentially overly conservative approach. To address the question of how these biased measures of interactions are suited for comparing modeled and empirical connectomes, we compared global model performance based on connectivity metrics that are sensitive and robust to zero-phase lags in this study. ICHOH, PLI, WPLI and LPC all showed a significantly lower match between simulated and empirical FC (around r=0.18) compared to coherence and PLV (Figure 8). For all six metrics, the global correlation was essentially abolished if the underlying SC was shuffled prior to simulation (yellow bars in (Figure 8)). Also, the overall model performance for ICOH, PLI, WPLI and LPC was considerably smaller than the mere correlation between SC and empirical FC (middle row in Figure 5A). What are the possible reasons for this performance drop with ICOH, PLI, WPLI and LPC? One reason could lie in the fact that the reference model SAR does not include delays, thus the simulated FC mainly consists of instantaneous interactions and a comparison with an empirical FC in which those interactions have largely been removed would be futile. However, the results were very similar using the Kuramoto model. The large-scale connectomes derived from all of the four biased metrics did not much reflect the coupling that emerged from our model of fast dynamics based on structural connectivity. Presumably, a considerable amount of functionally relevant synchrony takes place with near zero or zero-phase lag which is not detected using the biased scores. In fact, zero-phase lag synchronization has been detected between cortical regions in a visuomotor integration task in cats [88]. More recently, a study of spike train recordings showed how paths among somatosensory areas were dominated by instantaneous interactions [89]. But synchrony across areas incorporating delays can also lead to high coherence [90]. A recent modeling study investigated the detection rates of synchrony by different EEG phase synchronization measures (PLV, ICOH, WPLI) in a network of neural mass models. They found that no single phase synchronization measure performed substantially better than all the others, and PLV was the only metric able to detect phase interactions near ±0° or ±180° [81]. This study challenged the supposed superiority of biased metrics in practical applications, because they are biased against zero-phase interactions that do truly occur in the brain. Taken together we argue that by using biased metrics to detect neural synchrony a major portion of relevant coupling is neglected. However, as the relevant stage for comparisons is the source space, the undesired influence of volume conduction effects on the estimated connectivity is partly reduced [91]. Since effects of field spread can never be completely abolished also in the source space, we cannot rule out that volume conduction artifacts have influenced the high correlation in our model.

### 3.6 Conclusion

In summary, our framework demonstrates how technical alternatives and choices along the modeling path impact on the performance of a structurally informed computational model of global functional connectivity. We show that determining the resting-state alpha rhythm functional connectome, the anatomical skeleton has a major influence and that simulations of global network characteristics can further close the gap between brain network structure and function.

## Supporting Information

### S1 Empirical Data

#### Participants

Eighteen healthy volunteers (7 women, mean age 65.6±10.9 std) underwent DTI and EEG resting-state recording. None of the participants reported any history of serious medical, neurological or psychiatric diseases. None of the participants were taking any central nervous system-active medication. The study design was approved by the Local Ethical Committee of the Medical Association of Hamburg (PV 3777). All participants gave their written informed consent according to the ethical declaration of Helsinki.

#### MRI data acquisition

Structural imaging data were acquired using a 3 Tesla Siemens Skyra MRI scanner (Siemens, Erlangen, Germany) and a 32-channel head coil to acquire both diffusion-weighted and high-resolution T1-weighted anatomical images. For diffusion-weighted imaging, 75 axial slices were obtained covering the whole brain with gradients (b=1500 *mm*^2^/s) applied along 64 non-collinear directions with the sequence parameters: Repetition (TR) = 10000 ms, echo time (TE) = 82 ms, field of view (FOV) = 256x204, slice thickness (ST) = 2 mm, in-plane resolution (IPR) = 2x2 mm. The complete dataset consisted of 2 x 64 b1500 images and additionally one b0 image at the beginning and one after the first 64 images. For anatomical imaging, a three-dimensional magnetization-prepared, rapid acquisition gradient-echo sequence (MPRAGE) was used with the following parameters: TR = 2500 ms, TE = 2.12 ms, FOV = 240x192 mm, 256 axial slices, ST = 0.94 mm, IPR = 0.94 x 0.94 mm.

#### DTI data preprocessing and cortical parcellation

Diffusion-weighted images were analysed using the FSL software package 5.1 (http://www.fmrib.ox.ac.uk/fsl). All datasets were corrected for eddy currents and head motion. Fractional anisotropy (FA) maps were calculated fitting the diffusion tensor model at each voxel. Structural T1-weighted anatomical images were processed using the Freesurfer software package 5.3.0 with standard procedures and parameters resulting in a cortical parcellation of 68 cortical regions [92–94]. Accuracy of cortical parcellation was checked visually. Two homologous regions (left and right entorhinal cortex) were discarded from further analysis due to frequent imaging artefacts surrounding this area of the brain. The remaining set of 66 parcellated brain regions were used for further analysis. Registration of structural and diffusion images was achieved using linear and non-linear transformation tools implemented in FSL [95]. Each cortical parcellation was transformed to diffusion space using the non-linear transformation coefficient file and accuracy of registration checked individually.

#### Fiber tractography and structural connectome construction

Processing of diffusion data included application of a probabilistic diffusion model, modified to allow estimation of multiple (n=2) fiber directions using the program bedpostx [96, 97]. From each seed ROI voxel, 10000 samples were initiated through the probability distribution on principle fiber direction. Tracking resulted in individual maps representing the connectivity value between the seed ROI and individual voxels. Structural connectivity between two regions was measured masking each seed ROI results by each of the remaining ROI’s. In probabilistic tractography, connectivity distribution drops with distance from the seed mask. We calculated the average length between different ROI’s using the distance-correction option of probtrackx following recommendations from the online documentation of the FSL library. Values of average distances between seed and target ROI were applied alternatively to the reference method (Figure 1) to account for the confounding effect of tract length.

#### EEG data acquisition and analysis

Continuous EEG was recorded from 63 cephalic active surface electrodes arranged in the 10/10 system (actiCAP®, Brain Products GmbH, Gilching, Germany) during eight minutes eyes-open resting-state. Impedance was kept below 20 kΩ. Data were sampled at 1000 Hz, referenced to the Cz-electrode (actiCHamp^®^ amplifier, Brain Products GmbH, Gilching). One electrode was mounted below the left eye for EOG-recording. Electrode positions were registered using an ultrasound localization system (CMS20, Zebris, Isny, Germany) before EEG-recording. Patients were instructed to fixate a stationary fixation cross (viewangle ±5°) to reduce eye movements and were asked to avoid eye blinks, swallowing, any other movements and mental tasks like counting. The continuous EEG was offline rereferenced to a common cephalic average, demeaned, detrended and subjected to an independent component analysis (logistic infomax ICA; [98]) to remove eye-blink artifacts which were mostly reflected in 1-2 components. The data was downsampled to 125 Hz and segments containing artifacts like muscle activity, lead movements, electrode artifacts or incompletely rejected blink artifacts were removed visually.

The source activity was reconstructed using different inverse solutions:

1. an LCMV beamformer constrained by the covariance of the sensor data [44],
2. an ELORETA spatial filter [99] or
3. the MNE [69].

As a forward model we computed a boundary element method volume conduction model [65] based on individual T1-weighted structural MRI of the whole brain and individual electrode positions using the source space modeling functions of the SPM12 toolbox. The source time series were band pass filtered at the alpha frequency band (8±2 Hz) and a Hilbert transform applied. From there functional connectivity estimates were derived as explained above. Analysis was performed with the FieldTrip package for MEG/EEG data analysis [100] and the Statistical Parametric Mapping software (SPM12b, Wellcome Trust Centre for Neuroimaging, London, UK, http://www.fil.ion.ucl.ac.uk/spm) on MATLAB Version 7.12.0 (R2011a, The Mathworks Inc., Massachusetts, USA).

### S2 Frequencies

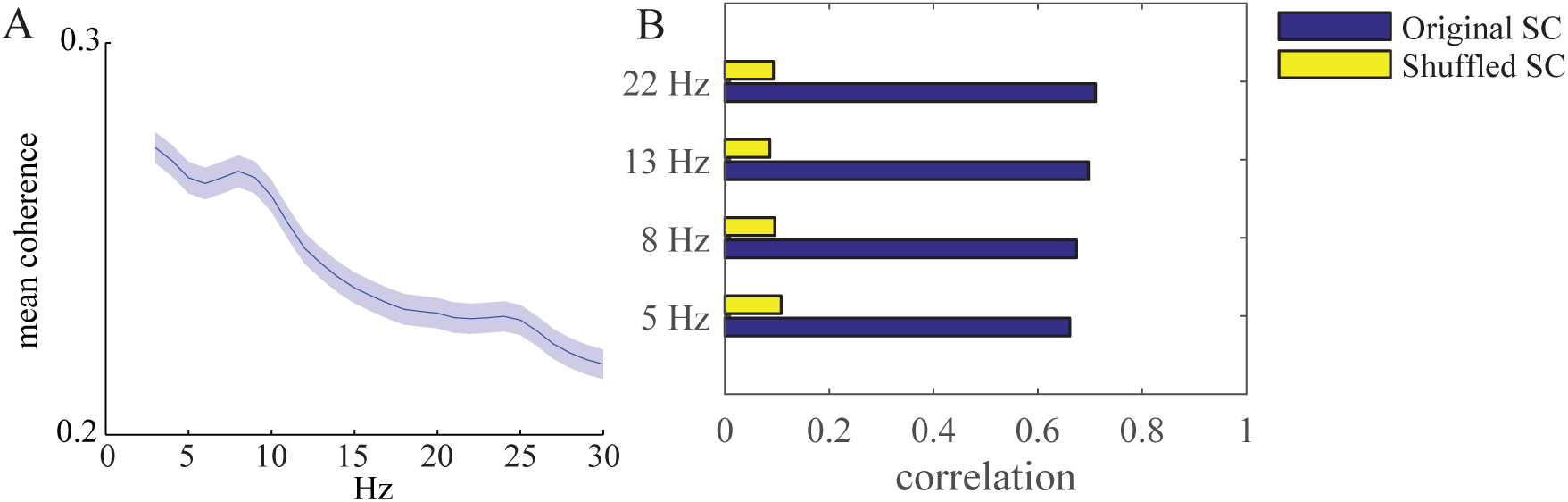
Evaluation of different EEG frequencies. A: The mean coherence values (±SEM, shaded area) between all ROIs (*n* = 2145) is calculated for the frequency range of 3-30 Hz. Overall coherence at lower frequencies is higher with a peak around 8 Hz and a smaller peak around 24 Hz. B: The model performance for different frequencies.

### S3 Connection strength and distance

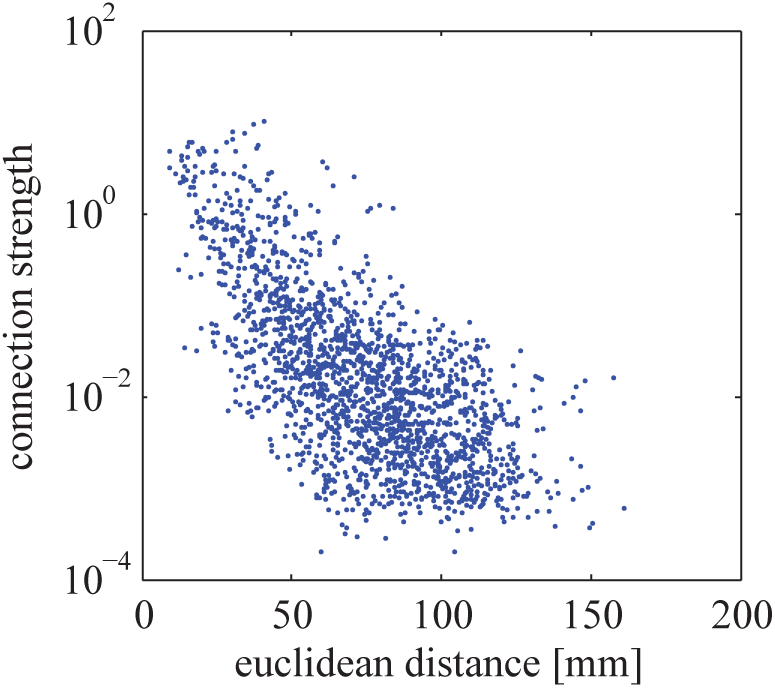
Dependence between connection strength and euclidean distance. The euclidean distance is measured between the center coordinates of individual ROIs. The strength between ROIs are the number of tracked DTI fibers divided by the product of both ROI sizes. The logarithm of the structural connection strength is inversely correlated with the euclidean distance (*r* = –0.37, *n* = 1883, *p* < .0001). Connections with zero strength (pairs of ROIs with no probabilistic tracked fibers between them) were excluded (n=262) due to the logarithmic axis.

## Acknowledgments

This work was funded by the DFG through SFB 936 Multi-Site Communication in the Brain. We would also like to thank Dr. Guido Nolte.

